# *FGF4* retrogene on CFA12 is responsible for chondrodystrophy and intervertebral disc disease in dogs

**DOI:** 10.1101/144022

**Authors:** Emily A. Brown, Peter J. Dickinson, Tamer Mansour, Beverly K. Sturges, Miriam Aguilar, Amy E. Young, Courtney Korff, Jenna Lind, Cassandra L. Ettinger, Samuel Varon, Rachel Pollard, C. Titus Brown, Terje Raudsepp, Danika L. Bannasch

**Affiliations:** Department of Population Health and Reproduction, School of Veterinary Medicine, University of California-Davis, Davis, CA 95616; Department of Surgical and Radiological Sciences, School of Veterinary Medicine, University of California-Davis, Davis, CA 95616; Department of Animal Science, University of California-Davis, Davis, CA 95616; Genome Center, University of California-Davis, Davis, CA 95616; Barney and Russum Animal Clinic, Fairfield, CA 94533; Department of Veterinary Integrative Biosciences, Texas A&M University, College Station, TX 77843

## Abstract

Chondrodystrophy in dogs is defined by dysplastic, shortened long bones and premature degeneration and calcification of intervertebral discs. Independent genome-wide association analyses for skeletal dysplasia (short limbs) within a single breed (p_Bonferroni_=0.0072) and intervertebral disc disease (IVDD) across breeds (p_Bonferroni_=4.02×10^−10^) both identified a significant association to the same region on CFA12. Whole genome sequencing identified a highly expressed *FGF4* retrogene within this shared region. The *FGF4* retrogene segregated with limb length and had an odds ratio of 51.23 (95% CI = 46.69, 56.20) for IVDD. Long bone length in dogs is a unique example of multiple disease-causing retrocopies of the same parental gene in a mammalian species. FGF signaling abnormalities have been associated with skeletal dysplasia in humans, and our findings present opportunities for both selective elimination of a medically and financially devastating disease in dogs and further understanding of the ever-growing complexity of retrogene biology.

---

Variation in domestic dog (*Canis familiaris*, CFA) morphology has long fascinated both scientists and pet owners. Domestication of the dog from the wolf and the subsequent variation in size and shape within purebred dog breeds is a remarkable feat of animal breeding and selection. One of the most extreme examples of dog breed differences is in limb length, as extremely short limbs define many breeds. This morphological feature is present in breeds from all over the world and from all American Kennel Club groups, indicating that the underlying genetic causes are likely very old.

Extensive examination of growth plates has been performed on many of these short-legged dog breeds (Dachshund, Pekingese, French Bulldog, Spaniels, Beagle), as these breeds are also prone to intervertebral disc disease (IVDD)^1–3^. Histopathological analysis of the bones of puppies from these breeds demonstrated that their short stature is due to defects in endochondral ossification, the process whereby cartilage is replaced with bone, in the developing limb. The long bone growth plates show disorganization of the proliferative zone and reduction in the depth of the maturation zone^1–4^. In addition to the long bones, similar but more subtle changes exist in endochondral ossification of the vertebral bodies^1,2^.

The intervertebral disc (IVD), which sits between vertebral bodies, is composed of an outer fibrous basket, called the annulus fibrosis, made of 70% collagen and an inner gel-like layer that is a remnant of the embryonic notochord, called the nucleus pulposus^5^. Together, these structures and the cartilaginous endplates allow for flexibility of the vertebral column. In chondrodystrophic dogs, the nucleus pulposus is gradually replaced by chondrocyte-like cells in chondroid metaplasia (or metamorphosis) that occurs between birth and 1 year of age^1,2^. Recent studies have shown that in advanced stages of degeneration in non-chondrodystrophoid dogs there is also replacement of notochordal cells by chondrocyte-like cells, similar to the changes observed in chondrodystrophoid dogs, although this happens at an older age^3,6–10^. The replacement of the nucleus pulposus with chondrocyte like cells is seen in humans, and chondrodystrophoid breeds have been proposed as models for human degenerative disc disease^3,7,11,12^.

Hansen described the two different types of canine IVD prolapse as type I and type II. Type I occurs exclusively in chondrodystrophic breeds and is characterized by premature degeneration of all discs in young dogs. In contrast, Type II occurs in older dogs and is usually limited to a single disc with only partial protrusion. In Type I disc disease the calcified nucleus pulposus may undergo an explosive herniation through the annulus fibrous into the vertebral canal, resulting in inflammation and hemorrhage and causing severe pain and neurological dysfunction^1,2^. In veterinary hospital population studies, breeds with a significant increased risk of IVDD include the Beagle, Cocker Spaniel, Dachshund, French Bulldog, Lhasa Apso, Pekingese, Pembroke Welsh Corgi, and Shih Tzu^13–15^. Pet insurance data suggests a conservative “lifetime prevalence” for IVDD in dogs of 3.5% in the overall population; however, in the chondrodystrophic breed with the highest risk, the Dachshund, the “lifetime prevalence” is between 20-62% with a mortality rate of 24%^9,16–19^. The effect of this disease on dogs and the financial burden to pet owners is enormous.

Skeletal dysplasia (SD), a general term to classify abnormalities of growth and development of cartilage and/or bone resulting in various forms of short stature, occurs in humans and dogs in many forms^20^. With advances in molecular genetics, many of the diseases in humans are being reclassified based on the specific underlying causative mutations^21^. To a lesser degree, progress has also been made in understanding the molecular nature of SD and the extreme interbreed limb length variation observed in dogs^22–25^. While the mutations causing some subtypes of SD in dogs have been determined, there are still many unexplained types of SD observed within and across dog breeds.

In 2009 the genetic basis for extreme differences in limb length in dogs was investigated by Parker et al. using an across breed genome-wide association approach^26^. They determined that a *FGF4* retrogene insertion on CFA18 was responsible for the “chondrodysplasia” phenotype in a number of breeds, such as the Basset Hound, Pembroke Welsh Corgi, and Dachshund. However, the *FGF4* retrogene insertion on CFA18 failed to explain breeds such as the American Cocker Spaniel, Beagle, and French Bulldog, that in addition to Dachshunds, were the breeds originally classified as chondrodystrophoid based on histopathologic and morphologic analysis by Hansen and Braund^1,3^. The FGF gene family has similarly been implicated in SD in humans, with mutations in *FGFR3* found to be responsible for achondrodysplasia, the most common form of dwarfism, characterized by shortened limbs and abnormal vertebrae and IVDs^21,27–31^. FGF genes are involved in a number of embryological development processes, and specific levels of ligand and receptor are key for appropriate growth and development^32–34^.

In this study, genome-wide association analysis in a cohort of Nova Scotia Duck Tolling Retrievers (NSDTRs) with and without severe SD identified a significant association on CFA12 due to a 12 Mb associated haplotype, of which 1.9 Mb was found to be shared in chondrodystrophoid breeds. Subsequent genome-wide association analysis of Hansen’s type I IVDD across breeds localized the same 1.9 Mb region on CFA12, suggesting that the locus responsible for SD in the NSDTR is also responsible for type I IVDD and the chondrodystrophoid phenotype across dog breeds. A previous genetic investigation of IVDD in Dachshunds and limb length morphology in Portuguese Water Dogs both identified the same CFA12 locus; however, neither study reported a causative mutation^35,36^. Here, using Illumina paired-end genomic sequencing, we uncover a second *FGF4* retrogene insertion (chr12:33.7 Mb (canFam3)) in the canine genome and show that it is not only responsible for SD in the NSDTR, but also chondrodystrophy, including the predisposition to Hansen’s type I IVDD, across all dog breeds.

## RESULTS

### GWAS for SD and IVDD

A form of SD is common in the NSDTR and is characterized by variable decrease in limb length and associated abnormalities, including long bone bowing, physeal widening, and joint incongruity. (Fig. 1a, 1b). On physical examination, in addition to shorter limbs, SD dogs may also have valgus limb deformities and larger ears (pinnae). While SD is a common phenotype in the breed, the degree of severity is highly variable.

**Fig. 1.**
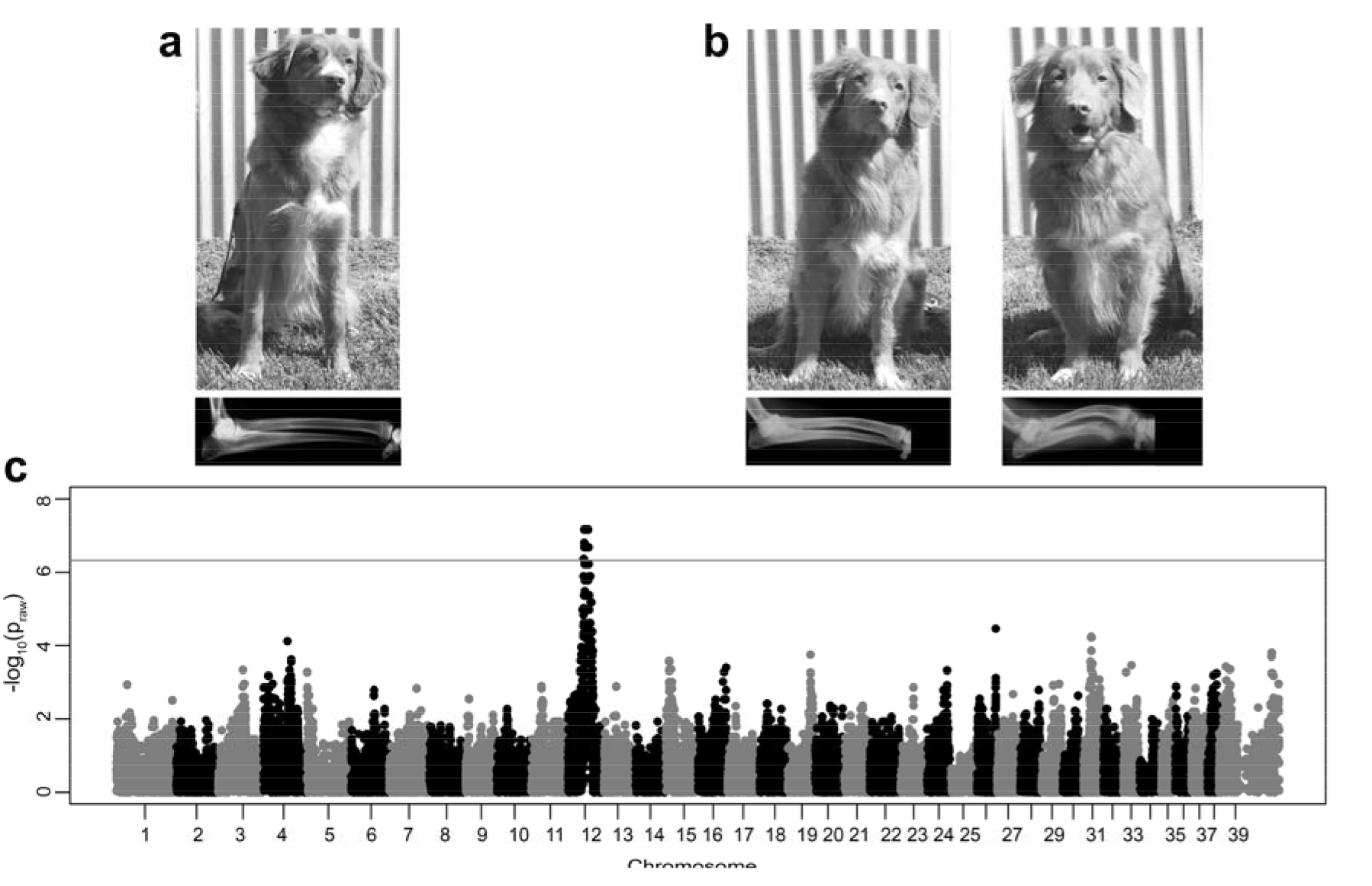
Skeletal Dysplasia in the NSDTR: a) Picture and lateral thoracic limb radiograph of unaffected NSDTRs (ages 1 year old and 4 years old, respectively). b) Left panels depict picture and lateral thoracic limb radiograph of mildly SD affected NSDTRs (ages 4 years old and 2 years old, respectively); right panels depict picture and lateral thoracic limb radiograph of more severely SD affected NSDTRs (ages 3 years old and 6 months old, respectively). Relative to the unaffected dog, the mildly SD affected NSDTR has cranial bowing of the radius. Radiographic changes in the more severely SD NSDTR include moderate cranial bowing of the radius, physeal widening, and incongruity of the elbow joint with the shape of the semi-lunar notch of the ulna being elongated. Pictures and radiographs are representatives of each phenotype and not paired (i.e. the radiographs are not of the dogs pictured). c) SD in NSDTR GWAS: Manhattan plot showing −log_10_ of the raw p-values for each genotyped SNP by chromosome (x-axis). After SNP quality control, there were 106,303 SNPs for Chi square analysis. Genomic inflation was 1.01604. Line denotes genome-wide significance based on Bonferroni corrected p-values.

To determine a region of the genome associated with SD in the NSDTR, genome-wide association analysis was performed using 13 NSDTR with severe SD and 15 NSDTR controls without severe SD. There were 41 SNPs that were genome-wide significant with a p_Bonferroni_<0.05, all present between chr12:35,413,695-46,117,273 (top SNP- chr12:36,790,324 p_Bonferroni_=0.007232) (canFam2) (Fig. 1c, Supplemental Fig. 1).

Underlying this strong association for SD in NSDTRs was an approximately 12 Mb critical interval from chr12:36-48 Mb (canFam2). Since the NSDTR SD phenotype is not uncommon in different dog breeds, we investigated haplotype sharing across breeds and observed that a portion of this associated haplotype was shared with two breeds of dog considered classically chondrodystrophic: the American Cocker Spaniel and Beagle^1,3^. By plotting the minor allele frequency (MAF) across this interval for 7 American Cocker Spaniels, 14 Beagles, and 13 SD affected NSDTR, the critical interval identified via GWAS for SD was shortened to a shared haplotype from chr12:36.4-38.3 Mb (canFam2) (Fig. 2a).

**Fig. 2.**
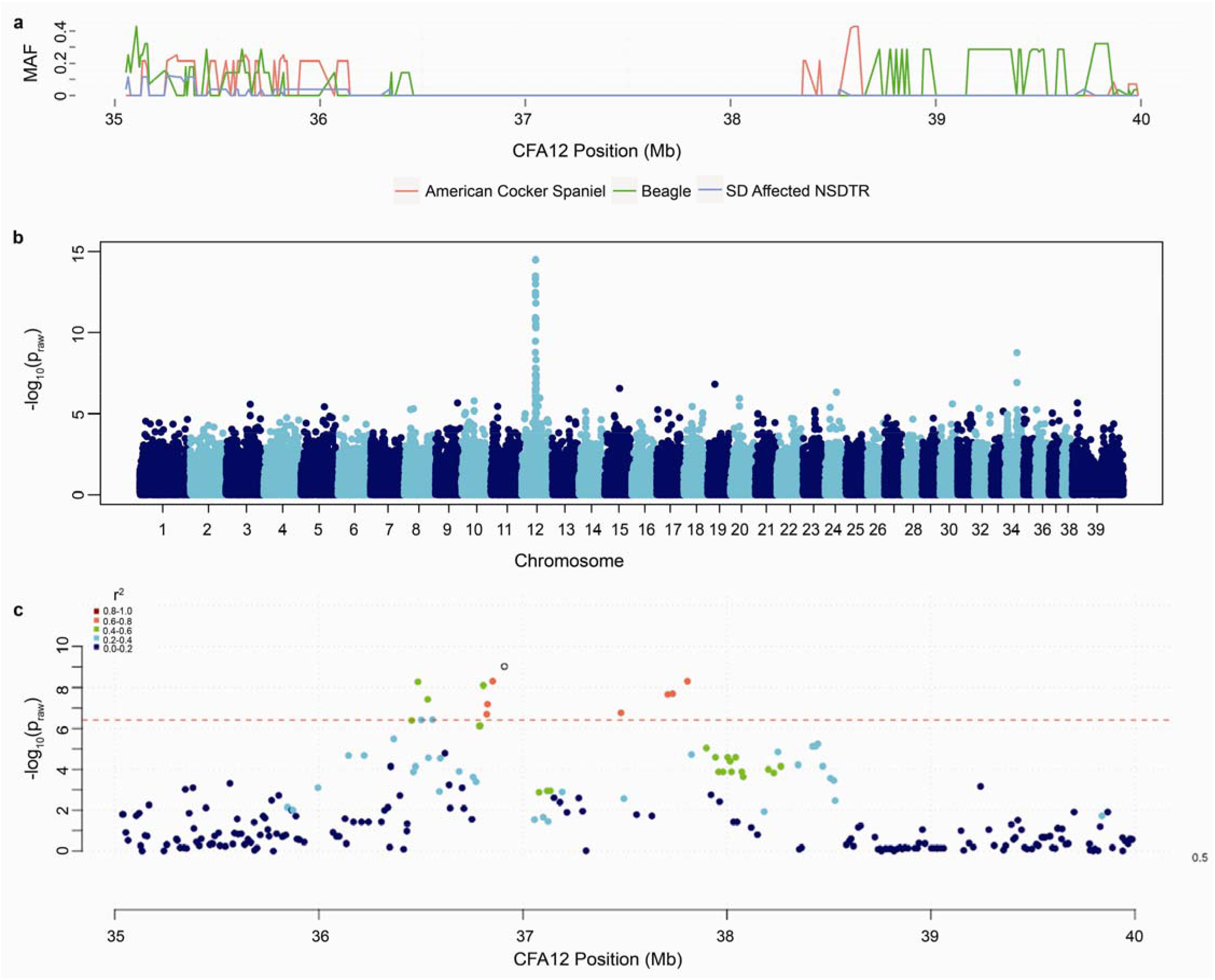
Across breed investigation of SD-IVDD locus: a) Minor allele frequency on the y-axis and base pair on CFA12 on the x-axis plotted by breed: SD affected NSDTR (n=13), American Cocker Spaniel (n=7), and Beagle (n=14). b) Manhattan plot for the SNPs in the across breed IVDD GWAS showing −log_10_ of the raw p-values (y-axis) for each genotyped SNP by chromosome (x-axis). After SNP quality control, there were 126,020 SNPs for Chi square analysis. Genomic inflation was 1.6339. c) SNPs in 5Mb region surrounding most highly associated SNP (chr12:36,909,311 (canFam2)) plotted by base pair on the x-axis and p-value on the y-axis. SNPs have been pruned from analysis using recommended criteria by Kierczak et al.^37^. SNPs are color-coded by r^2^ value to show the extent of linkage disequilibrium.

In order to test the hypothesis that the same locus was responsible for SD and chondrodystrophy, a second genome-wide association study was performed using IVDD affected cases (n=36) and unaffected controls (n=31) across 26 dog breeds (listed in Supplemental Table 6). The most highly associated SNP was located on CFA12 (chr12:36,909,311 (canFam2)) with a p_raw_=3.191x10^−15^, p_Bonferroni_=4.02×10^−10^, and odds ratio of 32.67 (Fig. 2b). Observing LD with the highest associated SNP using r^2^ values, the critical interval identified via GWAS for IVDD overlaps with that seen when mapping MAF across breeds and SD in the NSDTR (Fig. 2c).

### Identification of *FGF4* insertion on CFA12

To identify a causative variant for SD and IVDD, paired-end whole genome sequences of 2 cases, 1 SD affected NSDTR and 1 IVDD affected Dachshund, and 83 unaffected controls were investigated in the associated interval. There were 9,156 SNP variants and 7,877 insertion/deletion (indel) variants identified from chr12:33.1-35.5 Mb (canFam3) (chr12:36.1-38.5Mb (canFam2)); however, none segregated with the IVDD phenotype. The same interval was also investigated by visual inspection of BAM files to flag mate pairs with unusual insert sizes in an effort to identify any large indels. Using the 2 cases and 2 controls, 8 large indels (>200bp) were identified within the interval (Supplemental Table 1). 4 large indels did not segregate when investigated in additional control genomes, while the remaining 4 were eliminated after PCR showed lack of segregation between cases and controls.

Visual inspection of the BAM files for read-pairs mapping to a different chromosome location identified a region, located at approximately chr12:33,710,200 (canFam3), that segregated with the 2 cases and 2 controls (Supplemental Fig. 2). At this location, read mates mapped to chr18:48.4 Mb (canFam3) and chr7:68.3Mb (canFam3) in the NSDTR and Dachshund cases, but none of the controls. The reads that mapped to CFA18 aligned to endogenous *FGF4*, which was highly suggestive of a *FGF4* retrogene insertion at this location. The reads that mapped to CFA7 were investigated by PCR and appear to mark a genome assembly error or a mutation within the dog used for the genome assembly (canFam3).

To investigate the potential *FGF4* insert on CFA12, the region was PCR amplified using primers flanking the insertion site from genomic DNA of an IVDD affected Beagle. Wild type dogs without the insert had a single 615 bp band, while dogs homozygous for the CFA12 *FGF4* insertion had an approximately 4kb product. Sanger sequencing showed the insertion on CFA12 is 3,209 bp long (GenBank Accession # MF040221) and includes endogenous *FGF4* cDNA (i.e. *FGF4* exons spliced without introns), as shown in the insert schematic comparing endogenous *FGF4* to the CFA12 insert (Fig. 3). The insert also contains a majority of the predicted 5’UTR, which includes the transcription start site (TSS) as only PCR primers *FGF4_TSS.F1* and FGF4.R1 yielded a product in RT-PCR using cDNA from neonatal Beagle IVD (Supplemental Table 8).

**Fig. 3.**
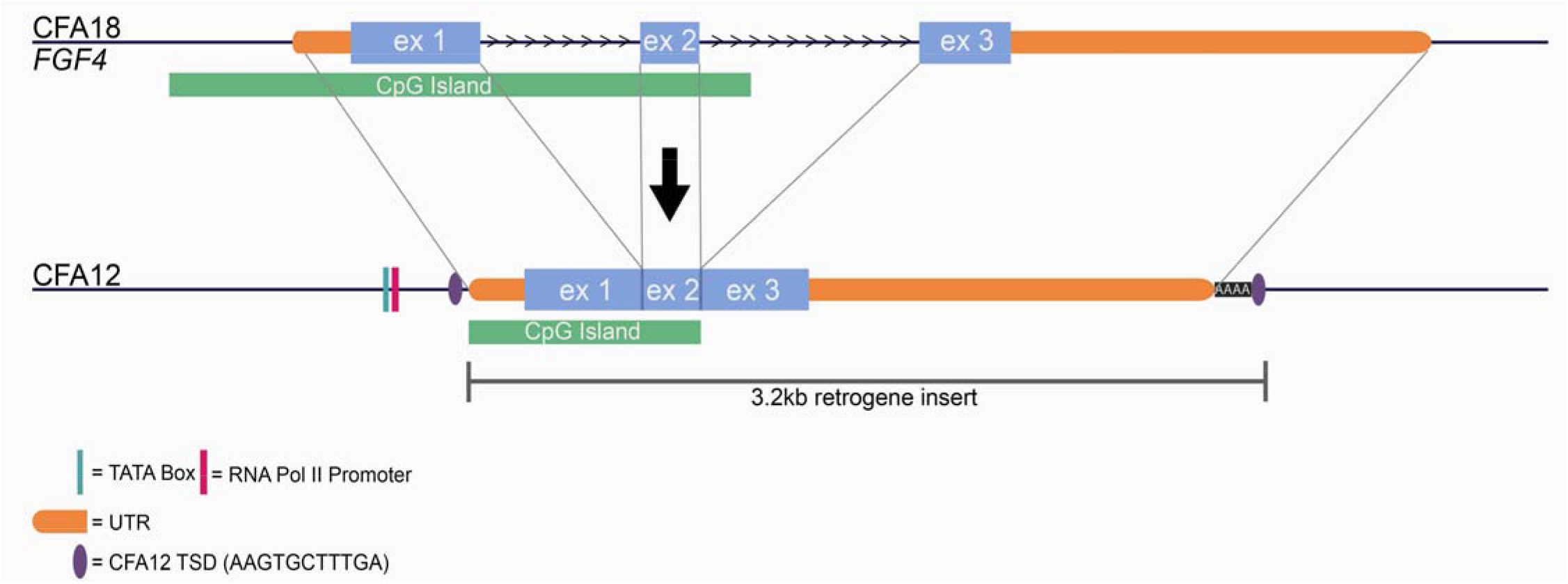
Schematic of endogenous FGF4 (CFA18) retrotransposition to CFA12:33,710,178 (canFam3): Predicted TATA box at chr12:33,709,940-947 (canFam3) and predicted RNA Pol II promoter at chr12:33,709,964-976 (canFam3), which are 239bp and 215 bp upstream, respectively^38^. Endogenous *FGF4* untranslated regions (UTR) are unknown in the dog; however, they were approximated in the figure based on human and mouse TransMap data available at the UCSC genome browser (genome.ucsc.edu). The predicted 5’UTR spans from chr18:48,413,185-48,413,480 (canFam3); however, RT-PCR in IVD from a Beagle suggest that the TSS is located between chr18:48,413,315-48,413,402 (canFam3). The insert includes all predicted 3’UTR, followed by 42 adenine residues and the duplicated 11bp TSD sequence (AAG TGC TTT GA) (chr12:33,710,168-33,710,178 (canFam3)). Endogenous FGF4 sequence that was retrotransposed also includes a large CpG island.

In order to compare the CFA12 *FGF4* retrogene to the previously identified CFA18 *FGF4* retrogene, it was necessary to obtain the full length sequence of the CFA18 insertion^26^. The cloned product was sequenced using the flanking and common internal primers (Supplemental Table 8), yielding a 2,665 bp insert (GenBank Accession # MF040222). While it contained the same length 5’UTR and *FGF4* cDNA as that seen in the CFA12 *FGF4* insert, the 3’UTR was shortened in comparison. The 3’UTR of the CFA18 *FGF4* insert was followed by a sequence containing 30 adenine and 1 guanine residues and a different target site duplication (TSD) sequence (AAG TCA GAC AGA G).

### Association of *FGF4* retrogene with SD, height, and IVDD

In order to assay the insertions in additional dogs, insertion and allele specific PCR based genotyping assays were developed for both the CFA12 *FGF4* insertion and the previously identified CFA18 *FGF4* insertion (Fig. 4a). Twelve SD NSDTR cases from the GWAS were genotyped and were homozygous for the CFA12 *FGF4* insertion, while all controls were heterozygous or wild type. Additionally, IVDD cases (n=7) from the NSDTR breed were collected and were either homozygous mutant or heterozygous for the CFA12 *FGF4* insertion (Supplemental Table 3). All NSDTR tested for the CFA18 *FGF4* insertion (n=31) were wild type, including SD and IVDD cases. NSDTRs with known height (n=20 males) at the withers were genotyped for the CFA12 *FGF4* insertion to investigate the association of height with genotype status. Height and genotype were significantly associated in a dose dependent manner when comparing wild type, heterozygous, and homozygous dogs (Fig. 4b).

**Fig. 4.**
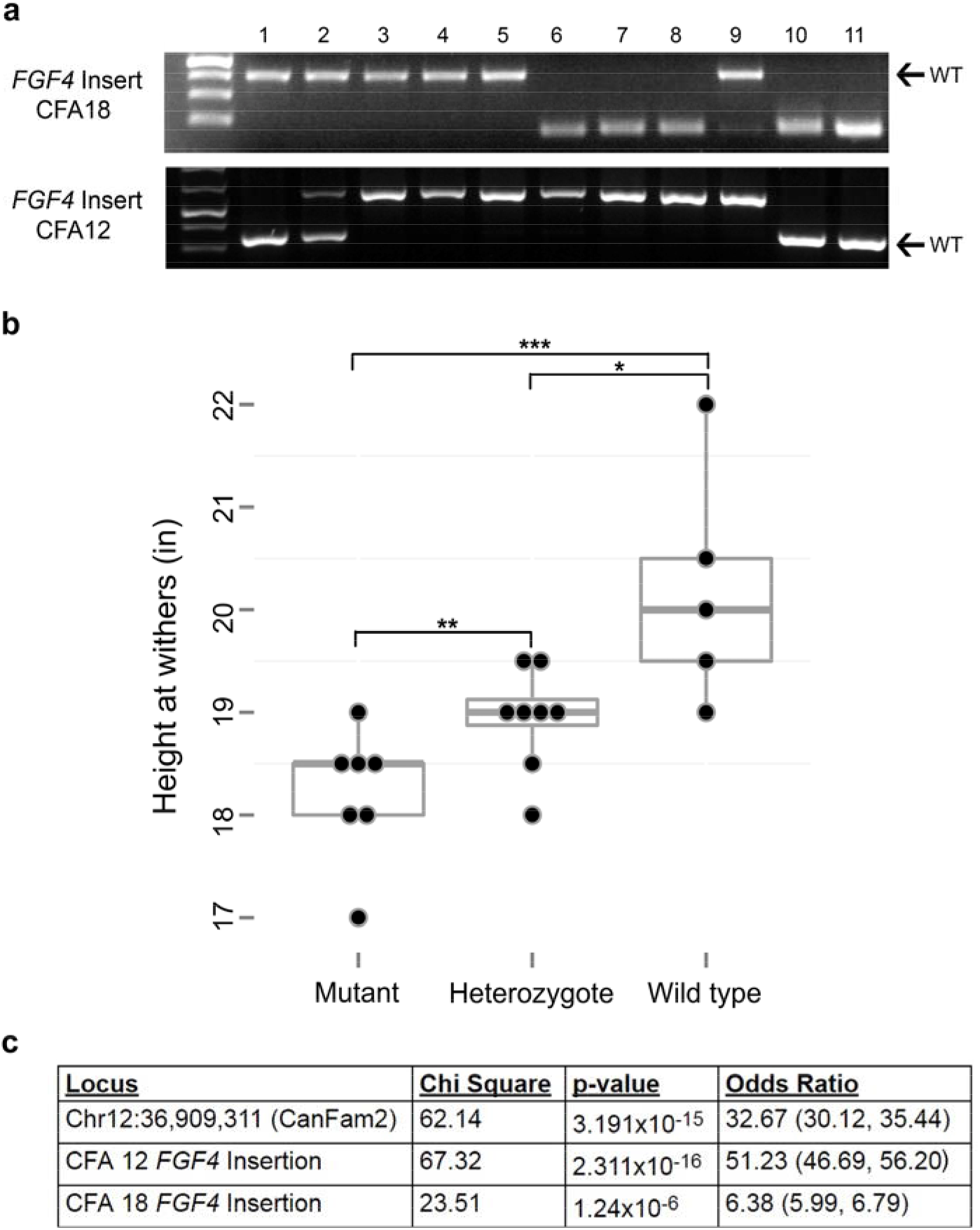
Association of *FGF4* insertion genotypes with height and IVDD: a) Genotyping results for CFA18 and CFA12 *FGF4* insertions across breeds. Arrows indicate wild type (WT) band. Lane order: Ladder; 1-3) NSDTR; 4) Beagle; 5) American Cocker Spaniel; 6) Dachshund; 7) Basset Hound; 8) Pembroke Welsh Corgi; 9) Coton de Tulear; 10) Cairn Terrier; 11) West Highland White Terrier. b) Height at the withers in inches (in) was available for 7 SD NSDTR cases: all were homozygous mutant for the CFA12 *FGF4* insertion, and their mean height was 18.22 in. Height was available for 13 NSDTR unaffected with SD: 5 were wild type and had a mean height of 20.2 in; 8 were heterozygous for the CFA12 *FGF4* insertion and had a mean height of 18.94 in. * indicate relative levels of association of the insertion with height. ***: p=0.007, **: p=0.016, *: p=0.034. c) Association of various identified loci with IVDD, including Chi square, p-value, and Odds ratio (95%confidence intervals in parenthesis).

To assess the significance of association of the CFA12 *FGF4* insertion with IVDD across breeds, dogs used in the IVDD GWAS were genotyped for both insertions. All dogs’ genotypes were concordant with phenotype except for one case, a Rottweiler (Supplemental Table 2). When associated with IVDD, the CFA12 *FGF4* insertion was more highly associated than both the most highly associated SNP from the GWAS, as well as the CFA18 *FGF4* insertion (Fig. 4c). To further investigate the association of the CFA12 *FGF4* insertion with IVDD, 33 additional cases were genotyped for the CFA12 *FGF4* insertion: 10 were heterozygous and 23 were homozygous for the CFA 12 *FGF4* insertion (Supplemental Table 3).

In order to investigate the breed distribution of the retrogene insertion, 568 dogs from 50 breeds were genotyped (Supplemental Table 4). The CFA12 *FGF4* insertion segregates in the majority of breeds where it occurs and is present in small and medium sized dog breeds with high frequency (Fig. 5). Interestingly, all of the dogs with the CFA12 *FGF4* insertion also have large external ears (pinnae), which is consistent with the phenotype seen in the NSDTR.

**Fig. 5.**
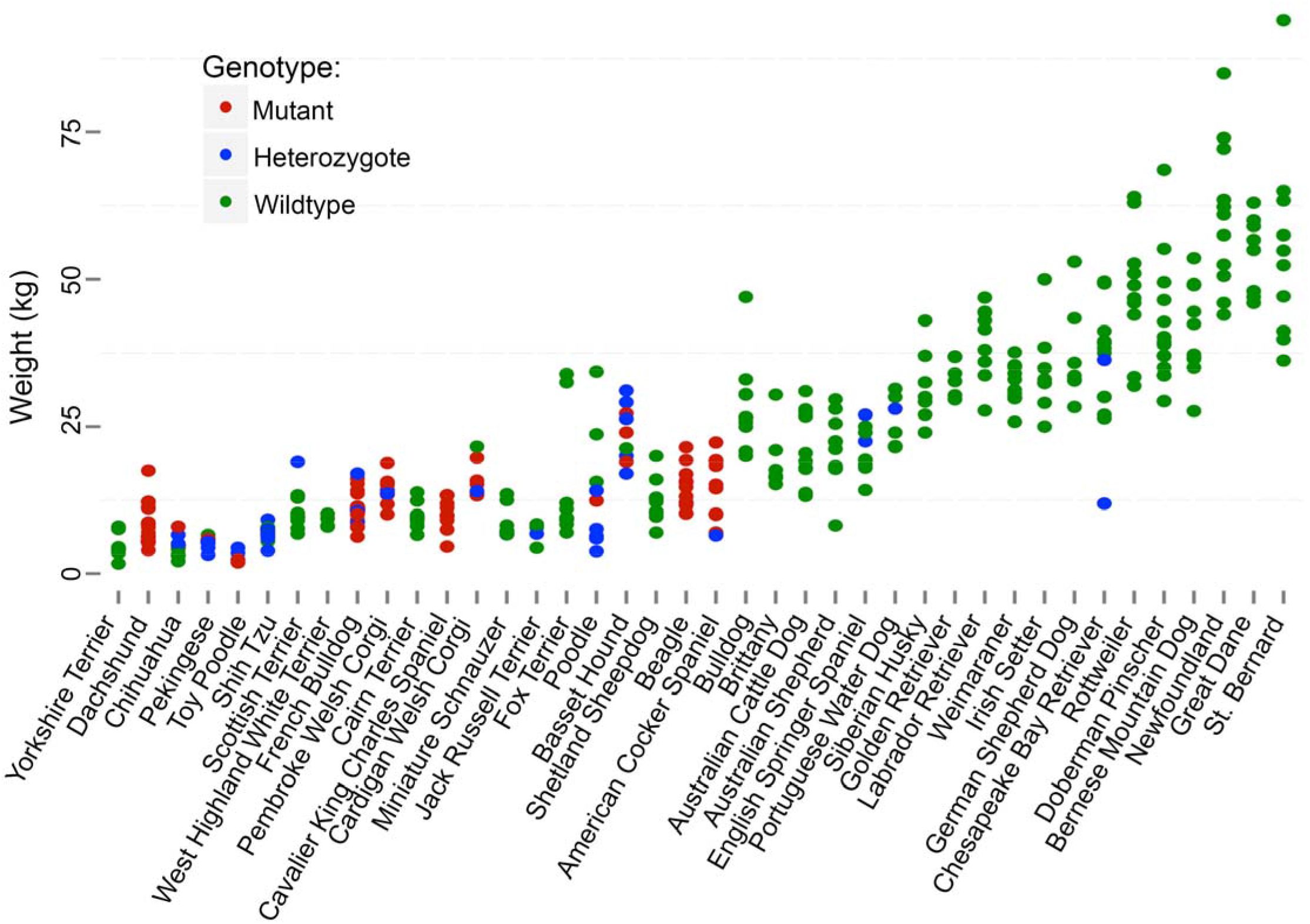
CFA 12 *FGF4* Genotypes Across Breeds: Genotypes for the CFA12 *FGF4* insertion across dog breeds ordered by breed standard height from shortest to tallest (x-axis), plotted by dog weight in kilograms (kg) (y-axis). Only genotyped dogs with weights available (n=376) were included in the figure. Dogs are color-coded by genotype status.

Based on occurrence of IVDD at the Pritchard Veterinary Medicine Teaching Hospital at UC Davis, the breeds with a statistically higher frequency of IVDD are also those with a higher frequency of the CFA12 *FGF4* insert allele, while the breeds with a statistically lower frequency of IVDD are those with a lower frequency of the CFA12 *FGF4* insert allele (Supplemental Table 5).

### CFA12 *FGF4* retrogene expression

To investigate the gene expression environment in which *FGF4* inserted on CFA12, semi-quantitative RT-PCR was performed for genes across the IVDD associated interval. Using cDNA derived from neonatal vertebral body (VB) and IVD, skeletal muscle, and testis, expression levels of genes across the CFA12 associated interval were assayed in a Beagle case and Cane Corso or Labrador Retriever control, including: *COL9A1, SMAP1, B3GAT2, OGFRL1, LINC00472, RIMS1, KCNQ5*, and *COL12A1*. Expression differences between case and control were not apparent in these genes; however, we confirmed that all except *RIMS1* are expressed in both neonatal VB and IVD, supporting that *FGF4* inserted itself in a gene milieu conducive to expression in IVD. Semi-q PCR for total *FGF4* (endogenous and retrogene products) in the same tissues showed increased expression across all tested tissue types in the case versus the control (Supplemental Fig. 3).

In order to evaluate the effect of the CFA12 *FGF4* retrogene insertion on overall *FGF4* transcript levels, quantitative RT-PCR was performed. A comparison between samples homozygous for the CFA12 *FGF4* insertion and samples with only the endogenous copy of *FGF4* (i.e. wild type for both the CFA12 and CFA18 *FGF4* insertions) showed a 19.47x higher (p=0.02857) and 2.16x higher (p=0.02857) expression of *FGF4* in neonatal IVD and VB, respectively (Supplemental Fig. 4).

## DISCUSSION

In this study, we report the identification of a *FGF4* retrogene insertion in the dog genome responsible for chondrodystrophy across dog breeds, characterized by both short limbs and susceptibility to Hansen’s type I intervertebral disc disease. A region was identified on CFA12 due to association with a segregating form of skeletal dysplasia observed in the NSDTR. While NSDTRs can be variably affected, the use of severely affected dogs enabled identification of the locus through GWAS. Haplotype sharing with chondrodystrophoid breeds and genome-wide association analysis for type I IVDD identified the same region on CFA12. Evaluation of mismapped mate pairs allowed the identification of a novel *FGF4* retrogene, which leads to an ~20 fold increase in expression of *FGF4* in neonatal intervertebral disc. Due to the embryonic expression pattern of *FGF4*, it is probable that these expression changes are also impacting endochondral ossification. This is the second *FGF4* retrogene identified in dogs that affects limb length. While the *FGF4* retrogene on CFA18 impacts limb length, the *FGF4* retrogene on CFA12 explains the chondrodystrophoid phenotype, which includes limb length and IVDD (significant odds ratio >50).

Fibroblast Growth Factor 4 (*FGF4*) is a growth factor gene expressed in specific tissues and at specific times throughout embryonic development in the mouse^39^. *FGF4* is highly expressed in the apical ectodermal ridge of the developing limb bud, as well as somites and the notochord that will form the vertebral column and IVDs^39–41^. FGF signaling is required for appropriate embryonic axial growth and segmentation, and *FGF4/FGF8* murine hypomorphs are characterized by altered vertebral morphology and smaller limb buds^42,43^. Additionally, *FGF8* hypomorphs are observed to have either hypoplastic or non-existent external ear structures^44^. In mice, creation of a gain of function *FGF4* copy to replace an inactive *FGF8* gene was able rescue limb development; however, it also caused abnormal tissue deposition and postaxial polydactyly, highlighting that levels of *FGF* throughout embryonic development must be properly controlled for normal limb formation^32^. While the specific embryonic expression pattern of *FGF4* in dogs with 4-6 copies of the gene is unknown, we hypothesize that the insertion site milieu on CFA12 versus CFA18 is contributing to differences in expression between the retrogenes, leading to the differences in phenotype.

A survey of retrogenes in the canine reference genome reported ~70 functional retrogenes in the dog; however, only the previous CFA18 *FGF4* retrogene insertion has been reported to be associated with a disease causing phenotype^26,45^. Similarly in humans, the formation of processed pseudogenes in general, as well as those that retain their intended function and cause disease, is rare^46–51^.

Both copies of the canine *FGF4* retrogenes have signatures of having arisen from RNA retrotransposed by LINE-1 integrase and reverse transcriptase, including flanking TSDs and polyA tracts (class 1 templated sequence insertion polymorphism)^52^. The CFA18 *FGF4* retrogene insertion was predicted to be expressed due to insertion near sequence with promoter properties^26^. While the CFA12 *FGF4* insertion is placed near a potential TATA box and RNA Pol II promoter, it is more likely that the CpG island included in the retrogene is driving expression^53–55^. This hypothesis is supported by the finding that a majority of retrogene expression is actually due to genomic context and contribution of CpG islands, not through the use of nearby promoters^56^. To our knowledge, this is the first documentation of a second retrogene insertion of the same parental gene resulting in a disease phenotype in a mammalian species. Due to the lack of resources available to identify these types of mutations, it is likely that there are other phenotype inducing retrocopies present in the canine genome that have yet to be discovered.

Chondrodystrophy associated mutation events occurred a very long time ago, as there are descriptions of short-legged dogs dating back over 4000 years^57^. In addition, both mutations occur concurrently in very unrelated dog breeds from diverse breed groupings and geographical locations. The fact that *FGF4* has been retrotransposed twice in dogs in the last 3-4 thousand years makes it likely that this has happened at other times. The large CpG island in the 5’ end of the endogenous *FGF4* gene may enable phenotypic consequences more readily than for other retrogenes. Once the *FGF4* retrogene appeared and produced an obvious phenotype, strong selection was likely applied to retain it, aided by the semi-dominant nature of the mutation.

The NSDTR is the smallest of the retriever dog breeds, and based on the association of the CFA12 *FGF4* insertion with height, we hypothesize that the heterozygous phenotype is aesthetically desirable and that selection is maintaining the insertion at a relatively high allele frequency. Investigation of the CFA12 *FGF4* insertion in additional breeds also showed high allele frequency in multiple small and medium sized dog breeds. In breeds also containing the CFA18 *FGF4* insertion, there is an even more dramatic decrease in height (e.g. Basset Hound, Cardigan Welsh Corgi, Dachshund, etc.), supporting that both *FGF4* retrogenes affect long bone length.

In addition to segregating with height, the CFA12 *FGF4* insertion also segregates with Hansen’s type I IVDD susceptibility. Of the IVDD cases genotyped for the CFA12 *FGF4* insertion, all were homozygous mutant or heterozygous, except for 1, suggesting that one additional copy of *FGF4* on CFA12 is sufficient to cause type I IVDD. The single discordant case was a Rottweiler, a breed that does not fit the chondrodystrophic phenotype. It is possible that there is another cause of IVDD in non-chondrodystrophoid dog breeds occurring without endochondral ossification defects^9^. IVDD-affected NSDTRs were also all either homozygous or heterozygous for the CFA12 *FGF4* insertion. This supports that while the CFA12 *FGF4* insertion is semi-dominant with respect to height, it is dominant for altered IVDs. Given that the CFA18 *FGF4* insertion is not found in the NSDTR and was inconsistently present in the IVDD cases tested, this further supports that the identified insertion on CFA12 is causing both short limbs and Hansen’s type I IVDD in both the NSDTR and across dog breeds.

The breeds with a higher frequency of the CFA12 *FGF4* insertion are the same breeds identified in the last 50 years as being predisposed to IVDD. Presence of the CFA18 *FGF4* insertion is common in many breeds with IVDD, and it is possible that it may contribute to the disease; however, previous mapping within Dachshunds, which are reported “fixed” for the CFA18 *FGF4* insertion, show segregation of the associated haplotype on chromosome 12 with IVDD, supporting that the CFA12 *FGF4* insertion is the critical factor determining disease status^26,35^. Of particular interest is the lack of reports of IVDD cases in breeds such as the Cairn Terrier and West Highland White Terrier, both of which have the CFA18 *FGF4* insertion, but not the CFA12 *FGF4* insertion. Similarly, the high incidence of IVDD in breeds such as the American Cocker Spaniel, Beagle, and French Bulldog that do not have the CFA18 *FGF4* insertion but a high frequency of the CFA12 *FGF4* insertion supports that *FGF4* specifically from CFA12 is contributing to the IVDD phenotype.

The segregation of the CFA12 *FGF4* insertion within dog breeds presents an opportunity for improvement of animal health, as implementation of genetic testing over time could lead to the elimination of type I IVDD. Based on the ever-growing popularity of some breeds, the number of animals with this intervertebral disc disease mutation across the globe is in the millions. Myelopathy secondary to IVD herniation is the most commonly presenting neurological disorder of the spinal cord in dogs^58^. The overall heath and financial consequences across the spectrum of presentations in companion dogs is immense. Prevention of disease through breeding and eradication has the potential for far-reaching benefits beyond those achievable through advances in surgical or medical therapy. Additionally, the dog may serve as a valuable human-animal model for IVDD. Administration of a tyrosine kinase inhibitor in a mouse model with a gain of function mutation in *FGFR3* has been shown to overcome growth defects associated with altered FGF signaling^33^. Based on the phenotype and molecular etiology of chondrodystrophy and IVDD in dogs, it has the potential to serve as a bridge between mouse and human studies evaluating the efficacy of targeted pharmacological treatment of FGF based genetic disorders.

Given the high mortality rate of IVDD and the high cost of surgery, identification of this susceptibility locus could provide a valuable tool for owners, breeders, and veterinarians for mitigating risk of intervertebral disc herniation and resulting myelopathy^9^. This could be especially useful in breeds that have both the CFA12 and CFA18 *FGF4* retrogene, as they could breed away from the CFA12 *FGF4* retrogene, while still maintaining the aesthetically desirable shortness in stature contributed by the CFA18 *FGF4* retrogene. In breeds with only the CFA12 *FGF4* retrogene, breeders will ultimately decide if prevention of Hansen’s type I IVDD outweighs any potential loss of shortness (or gain in height).

## METHODS

Methods are available in the online version of the paper.

## ACKNOWLEDGEMENTS

Maxine Adler Endowed Chair Funds

Center for Companion Animal Health, School of Veterinary Medicine, University of California, Davis NIH NIDCR R01DE22532

UC Davis Signature Research in Genomics (SRG): ACTG

NIH T35 OD010956

NIH T32 OD010931

## AUTHOR CONTRIBUTIONS

DLB designed the experiments. DLB, EAB, PJD, CK, JL, CLE, AEY, MA, TM, TB, TR performed experiments. DLB, EAB, PJD, JL, AEY, MA, SV collected samples. RP interpreted canine radiographs. DLB and EAB analyzed data. EAB, DLB, and PJD wrote the manuscript with input from all other authors.

## ONLINE METHODS

### Phenotype and Sample Collection

Blood samples, height at the withers, thoracic limb radiographs, and pictures (when possible), were collected from privately owned NSDTRs affected with owner or veterinarian reported skeletal dysplasia (SD), as well as phenotypically “normal” dogs. Additionally, blood samples were collected from cases of type I intervertebral disc disease (IVDD) seen at the University of California, Davis School of Veterinary Medicine Teaching Hospital and privately owned NSDTRs. IVDD cases were defined by the presence of one or more mineralized thoracolumbar intervertebral discs (IVD), as confirmed by vertebral column radiographs and/or the presence of extruded calcified degenerative disc material at surgery or necropsy. DNA was extracted from EDTA whole blood samples using Gentra Puregene DNA purification extraction kit (Qiagen, Valencia, CA). Collection of canine samples was approved by the University of California, Davis Animal Care and Use Committee (protocol #18561).

### Genome-wide Association Study

Genome-wide single nucleotide polymorphism (SNP) genotyping was performed using the Illumina Canine HD 174,000 SNP array (Illumina, San Diego, CA) for 13 NSDTR SD cases and 15 NSDTR controls with no reported SD. SNPs were pruned from analysis if the minor allele frequency was <5% and the call rate <90%. Additionally, a separate GWAS was performed for 36 IVDD cases from 16 breeds and 31 controls with no reported IVDD from 14 breeds (number of dogs from each breed listed in Supplemental Table 6). The SNPs were pruned from analysis if the minor allele frequency was <10% and the call rate <90%. Chi-square association analysis, Bonferroni adjustments, and genomic inflation calculations were performed in Plink^59^. Figures were made in R using ggplot2 and cgmisc packages^37,60^.

### WGS

For library prep and sequencing, DNA was fragmented using the Covaris E220 sonicator (Covaris Inc.), and then followed by selection of 550 bp insert size fragments. Illumina paired-end 150 bp libraries were prepared using PCR-free library prep kits. Sequencing was done on the HiSeq2500 platform at the BGI sequencing facility. Reads were scanned for sequencing adaptors and low quality sequences using the Trimmomatic software package (V 0.36)^61^. High quality reads were aligned to the dog reference genome canFam3^62^ using the BWA-MEM algorithm of the BWA software package (v0.7.7)^63^. Duplicate reads were excluded using Picard v2.2.4 tool MarkDuplicates^64^. Variant calling was performed using GATK HaplotypeCaller (v3.5)^65^. SNPs and small insertions and deletions (indels) were investigated for segregation with the IVDD phenotype in a 4Mb region (chr12:33.1-35.5 Mb (canFam3)), which included the critical interval identified using GWAS. Segregation of variants was performed using 2 cases (1 affected NSDTR and 1 Dachshund) compared to 83 controls of various normal legged breeds. To investigate the presence of large indels within the critical interval, BAM files covering the associated interval were scanned by eye using the Integrative Genomics Viewer (IGV, Broad Institute) in 2 cases (1 Dachshund and 1 SD affected NSDTR) and 2 controls (1 NSDTR and 1 Saluki) for segregation with the IVDD phenotype. The reads were viewed as color-coded by insert size and pair orientation to flag mate pairs that mapped to other chromosomes in order to easily identify large insertions and deletions.

### Investigation of Large Indels

BAM files for additional control genomes (1 NSDTR, 1 Weimaraner, 1 Border Collie (https://www.ncbi.nlm.nih.gov/biosample/SAMN03801652)) were used to evaluate segregation of the 8 identified large indels. The remaining segregating large indels (Deletions 1-3 and Insert 5) were investigated in additional cases and controls using PCR (primers listed in Supplemental Table 7).

### Investigation of potential insert from CFA7

The integrity of the region on CFA7 (approximately chr7:68,371,500-68,374,000) was tested using PCR and sequencing, as read mates at the *FGF4* insertion site on CFA12 also mapped to this location. Primers (listed in Supplemental Table 7) spanning the potentially inserted segment of CFA7 were used in PCR for 6 cases (1 NSDTR, 1 Beagle, 1 Basset Hound, 1 French Bulldog, and 1 Maltese) and 1 control (Boston Terrier) using the LongAmp Taq PCR Kit (New England Biolabs, Ipswich, MA, USA). If the genome assembly is correct, PCR product size should be 2,077bp, if not, the product would only be 842bp. PCR products for 3 cases and 1 control were sequenced on an Applied Biosystems 3500 Genetic Analyzer using the Big Dye Terminator Sequencing Kit (Life Technologies, Burlington, ON, Canada) and the products aligned to the UCSC genome browser using BLAT (genome.ucsc.edu/).

### Cloning

To obtain the full *FGF4* retrogene insertion sequence on CFA12, as well as on CFA18 for comparison, the PCR products using CFA12 *FGF4* Insertion and CFA18 *FGF4* Insertion primer pairs (Supplemental Table 8), respectively, were cloned using the TOPO TA Cloning kit with PCR2.1 TOPO (Thermo Fisher Scientific, Inc., Waltham, MA, USA) and One Shot TOP10 competent cells. CFA12 and 18 *FGF4* insertions were amplified from genomic DNA using the LongAmp Taq PCR Kit (New England Biolabs, Ipswich, MA, USA) using primers flanking the inserts on CFA12 and 18, respectively and recommended cycling conditions. The CFA12 *FGF4* insertion was cloned from a Beagle and the CFA18 *FGF4* insertion was cloned from a Dachshund. Plasmid DNA was extracted using the QIAprep Spin Miniprep Kit (Qiagen, Valencia, CA, USA). To confirm successful transformation, plasmid DNA was sequenced using vector primers M13.F and M13.R (Thermo Fisher Scientific, Inc., Waltham, MA, USA) on an Applied Biosystems 3500 Genetic Analyzer using the Big Dye Terminator Sequencing Kit (Life Technologies, Burlington, ON, Canada) and analyzed using VectorNTI software (Thermo Fisher Scientific, Inc., Waltham, MA, USA). Internal *FGF4* primers were also used to create overlapping contigs and ensure the entire insert was sequenced (listed in Supplemental Table 8). CFA12 and 18 *FGF4* inserts were aligned to endogenous *FGF4* sequence to confirm presence of the gene’s entire coding sequence. Polymorphisms identified were not queried in additional dogs, so could be due to sequencing or cloning error or be dog/breed specific.

### Transcription start site (TSS) Investigation

To ensure the CFA12 *FGF4* insertion included the transcription start site (TSS), PCR was performed using cDNA with primers at varying positions 5’ to the *FGF4* start codon (Supplemental Table 8). cDNA was synthesized from RNA extracted from neonatal intervertebral disc (IVD) and vertebral body (VB), as described below, from a Beagle. Amplified products were visualized on a 2% agarose gel.

### Genotyping Assay

The presence or absence of the *FGF4* insertion on CFA12 and 18 was assayed using a PCR-based genotyping test. Three primer PCR was performed using a forward and reverse primer flanking the respective insert, as well an additional forward primer located within the *FGF4* insert (listed in Supplemental Table 9). For the CFA12 *FGF4* insert assay, each reaction included 12μl water, 2μl 10x Buffer, 0.8μl 25mM MgCl_2_, 2μl dNTP, 0.5μl of the external forward primer (20μM), 0.7μl of the internal forward primer (20μM), 0.8μl of the reverse primer (20μM), 0.2μl of HotStarTaq DNA Polymerase (Qiagen, Valenica, CA, USA), and 1μl of DNA. For the CFA18 *FGF4* insert assay, each reaction included 12μl water, 2μl 10x Buffer, 0.8μl 25mM MgCl_2_, 2μl dNTP, 0.5μl of the external forward primer (20μM), 0.6μl of the internal forward primer (20μM), 0.9μl of the reverse primer (20μM), 0.2μl of HotStarTaq DNA Polymerase (Qiagen, Valenica, CA, USA), and 1μl of DNA. Amplified products were visualized on a 2% agarose gel. For the CFA18 *FGF4* insertion: wild type dogs had a single 388bp band, homozygous mutant samples had a single 168bp band, and both bands were present in heterozygous samples. For the CFA12 *FGF4* insertion, a single 333bp band was present in wild type samples, a single 654bp band was present in the homozygous mutant samples, and both bands were present in heterozygous samples.

Height at the withers was collected for 20 male NSDTRs to associate dog height with the CFA12 *FGF4* insertion. Significance was determined using a one-tailed T-test and a threshold cutoff of p<0.05.

To compare the significance of association of the CFA12 *FGF4* insertion to the most highly associated SNP and the CFA18 *FGF4* insertion, 34 of 36 IVDD cases and 31 controls were genotyped for both the CFA12 and CFA18 *FGF4* insertions. Chi square and odds ratio analysis was performed in Plink^59^.

### IVDD Incidence Across Breeds seen at UC Davis SVM VMTH

To assess frequency of IVDD in specific breeds, UC Davis School of Veterinary Medicine Teaching Hospital (VMTH) records were searched between 1980 and 2016 for dogs with a clinical diagnosis of “disc/k disease” or “IVDD.” Significantly over or underrepresented breeds relative to the total VMTH hospital population were determined based on a Chi-squared test. Significance was set at p<0.05.

### qRT-PCR

cDNA was prepared from RNA that was extracted from IVD or VB dissected from neonatal canine tail samples, skeletal muscle, and testis, using the QuantiTect Reverse Transcription Kit (Qiagen, Valencia, CA, USA) and the RNeasy Fibrous Tissue Mini Kit (Qiagen, Valencia, CA, USA), respectively. Additionally, cDNA was made from commercially available Beagle skeletal muscle and testis RNA (Zyagen, San Diego, CA, USA).

Semi-quantitative RT-PCR was performed for genes within and near the critical interval, including *COL9A1, SMAP1, B3GAT2, OGFRL1, LINC00472, RIMS1, KCNQ5*, and *COL12A1*, as well as *FGF4*, for VB, IVD, skeletal muscle, and testis cDNA from a case and control. *RPS5* was also included as a housekeeping gene control. All case samples were collected from a Beagle, while control VB and IVD were from a Cane Corso and skeletal muscle and testis from a Labrador Retriever. Primers spanning at least 1 intron were designed for all genes using Primer3, except for *RPS5* in which the primers were as recommended by Brinkhof et al.^66,67^. Each reaction included 13.9μl water, 2μl 10x Buffer with MgCl_2_, 1μl dNTP, 1μl of each forward and reverse primers (20μM) (listed in Supplemental Table 10), 1μl of HotStarTaq DNA Polymerase (Qiagen, Valenica, CA, USA), and 1μl of cDNA made from 1000ng of RNA. Amplified products were visualized on a 2% agarose gel.

Quantitative RT-PCR was performed for *FGF4* using cDNA synthesized from 500ng of RNA extracted from IVD and VB dissected from the tails of 4 cases (4 Beagle) and 5 controls (1 Rottweiler and 4 Cane Corso). A 2-step cycle protocol was employed using the Rotor-Gene SYBR Green PCR Kit (Qiagen, Valencia, CA, USA) on the Rotor Gene Q real-time PCR system: Initial denaturation at 95°C for 5 minutes; Annealing at 95°C for 5 seconds and extension at 60°C for 10 seconds for 35 cycles; Final melt curve. Samples were run in triplicate with 20ng of template cDNA each for both the IVD and VB experiment. *FGF4* transcript levels were normalized to *RPS5* and analyzed for fold change differences in expression using ΔΔC_T_. A technical replicate was removed from analysis if the standard deviation of the 3 technical replicates was greater than 1, and a sample was removed from analysis if there were less than 2 technical replicates that met this criteria. Fold change in expression of *FGF4* in IVD and VB was calculated by taking 2^−(ΔΔCT)^ for each tissue, respectively. Statistical significance was assessed using a Mann-Whitney-Wilcoxin test.

